# Prediction of prophages and their host ranges in pathogenic and commensal *Neisseria* species

**DOI:** 10.1101/2021.12.16.473053

**Authors:** Giulia Orazi, Alan J. Collins, Rachel J. Whitaker

**Author notes:** **Corresponding authors** Giulia Orazi, Rachel J. Whitaker.

## Abstract

The genus *Neisseria* includes two pathogenic species, *N. gonorrhoeae* and *N. meningitidis*, and numerous commensal species. *Neisseria* species frequently exchange DNA with one other, primarily via transformation and homologous recombination, and via multiple types of mobile genetic elements (MGEs). Few *Neisseria* bacteriophages (phages) have been identified and their impact on bacterial physiology is poorly understood. Furthermore, little is known about the range of species that *Neisseria* phages can infect. In this study, we used three virus prediction tools to scan 248 genomes of 21 different *Neisseria* species and identified 1302 unique predicted prophages. Using comparative genomics, we found that many predictions are dissimilar from other prophages and MGEs previously described to infect *Neisseria* species. We also identified similar predicted prophages in genomes of different *Neisseria* species. Additionally, we examined CRISPR-Cas targeting of each *Neisseria* genome and predicted prophage. While CRISPR targeting of chromosomal DNA appears to be common among several *Neisseria* species, we found that 20% of the prophages we predicted are targeted significantly more than the rest of the bacterial genome in which they were identified (i.e., backbone). Furthermore, many predicted prophages are targeted by CRISPR spacers encoded by other species. We then used these results to infer additional host species of known *Neisseria* prophages and predictions that are highly targeted relative to the backbone. Together, our results suggest that we have identified novel *Neisseria* prophages, several of which may infect multiple *Neisseria* species. These findings have important implications for understanding horizontal gene transfer between members of this genus.

**IMPORTANCE:** Drug-resistant *Neisseria gonorrhoeae* is a major threat to human health. Commensal *Neisseria* species are thought to serve as reservoirs of antibiotic resistance and virulence genes for the pathogenic species *N. gonorrhoeae* and *N. meningitidis*. Therefore, it is important to understand both the diversity of mobile genetic elements (MGEs) that can mediate horizontal gene transfer within this genus, and the breadth of species these MGEs can infect. In particular, few bacteriophages (phages) have been identified and characterized in *Neisseria* species. In this study, we identified a large number of candidate phages integrated within the genomes of commensal and pathogenic *Neisseria* species, many of which appear to be novel phages. Importantly, we discovered extensive interspecies targeting of predicted phages by *Neisseria* CRISPR-Cas systems, which may reflect their movement between different species. Uncovering the diversity and host range of phages is essential for understanding how they influence the evolution of their microbial hosts.

## INTRODUCTION

The genus *Neisseria* includes the human pathogens *N. gonorrhoeae* and *N. meningitidis* as well as a multitude of highly diverse commensal species that colonize mucosal surfaces of humans and animals (1). Because of the extensive spread of antibiotic resistance (AR) among strains of *N. gonorrhoeae*, infections caused by this pathogen are becoming increasingly difficult to treat (2). Consequently, the WHO and CDC consider *N. gonorrhoeae* a high-priority and urgent threat among antibiotic-resistant pathogens (3, 4).

*Neisseria* species are naturally competent and frequently exchange DNA with one other via transformation and homologous recombination (5–8). MGEs, such as plasmids, genetic islands, and bacteriophages (phages) can also mobilize genetic material and are powerful forces in shaping bacterial evolution (9–14). Phages are incredibly abundant and can deeply influence the fitness and virulence of their bacterial hosts, particularly when integrated into the bacterial chromosome (15–20). For example, the filamentous prophage MDAΦ promotes attachment of *N. meningitidis* to epithelial cell monolayers (18) and is associated with the ability of this pathogen to cause invasive disease (17).

In contrast to many highly studied Gammaproteobacteria prophages, few have been identified and characterized in the Betaproteobacteria (21–23). *Neisseria* prophages have been identified primarily in *N. gonorrhoeae* and *N. meningitidis* and consist of a small number of filamentous (24–27) and double-stranded DNA (dsDNA) prophages (28), including Mu-like prophages (21, 29–32). With the exception of MDAΦ, the impact of phages on *Neisseria* biology and pathogenicity remains poorly understood (13). Furthermore, few studies have investigated the host ranges of *Neisseria* phages (33, 34).

Microbes can defend themselves against phages and other MGEs using a variety of systems. One such system is CRISPR-Cas, which is composed of Clustered Regularly Interspaced Short Palindromic Repeats (CRISPR) arrays and CRISPR-associated (Cas) proteins. Importantly, sequence identity between the spacer and the MGE is required for immunity, which means that CRISPR arrays contain a record of previous encounters with MGEs within the sequences of their spacers. Therefore, this historical record can be used to infer the bacterial hosts of viruses (35–39). Approximately 40% of *N. meningitidis* genomes encode Type II-C CRISPR arrays (40, 41), and multiple putative CRISPR systems have been identified in several commensal species (41–43). In contrast, only degenerate CRISPR arrays have been found in *N. gonorrhoeae* (41, 42).

In this study, we sought to uncover novel *Neisseria* phage diversity. We used bioinformatic virus prediction tools to scan publicly available genomes of pathogenic and commensal *Neisseria* species for prophages. Using comparative genomics, we found that many of these predictions are dissimilar from previously identified *Neisseria* MGEs and are potential targets of CRISPR-Cas systems. Finally, we used interspecies CRISPR targeting of known and predicted prophages to infer whether they may infect multiple different *Neisseria* species.

## RESULTS

### Predicting prophages within genomes of pathogenic and commensal *Neisseria* species

To search for prophages, we compiled a dataset of 248 publicly available high-quality *Neisseria* genome assemblies from GenBank (44, 45) that includes *N. gonorrhoeae, N. meningitidis*, and 19 commensal species (see Methods and **Table S1A** for a description of the dataset). The relationships between the genomes in this dataset are shown in a phylogenetic tree based on ribosomal gene sequences and a heatmap of the average nucleotide identity (ANI) between each pair of genomes (**Fig. 1**). The phylogeny presented here is consistent with previously reported relationships between *Neisseria* species (46).

**Fig. 1.**
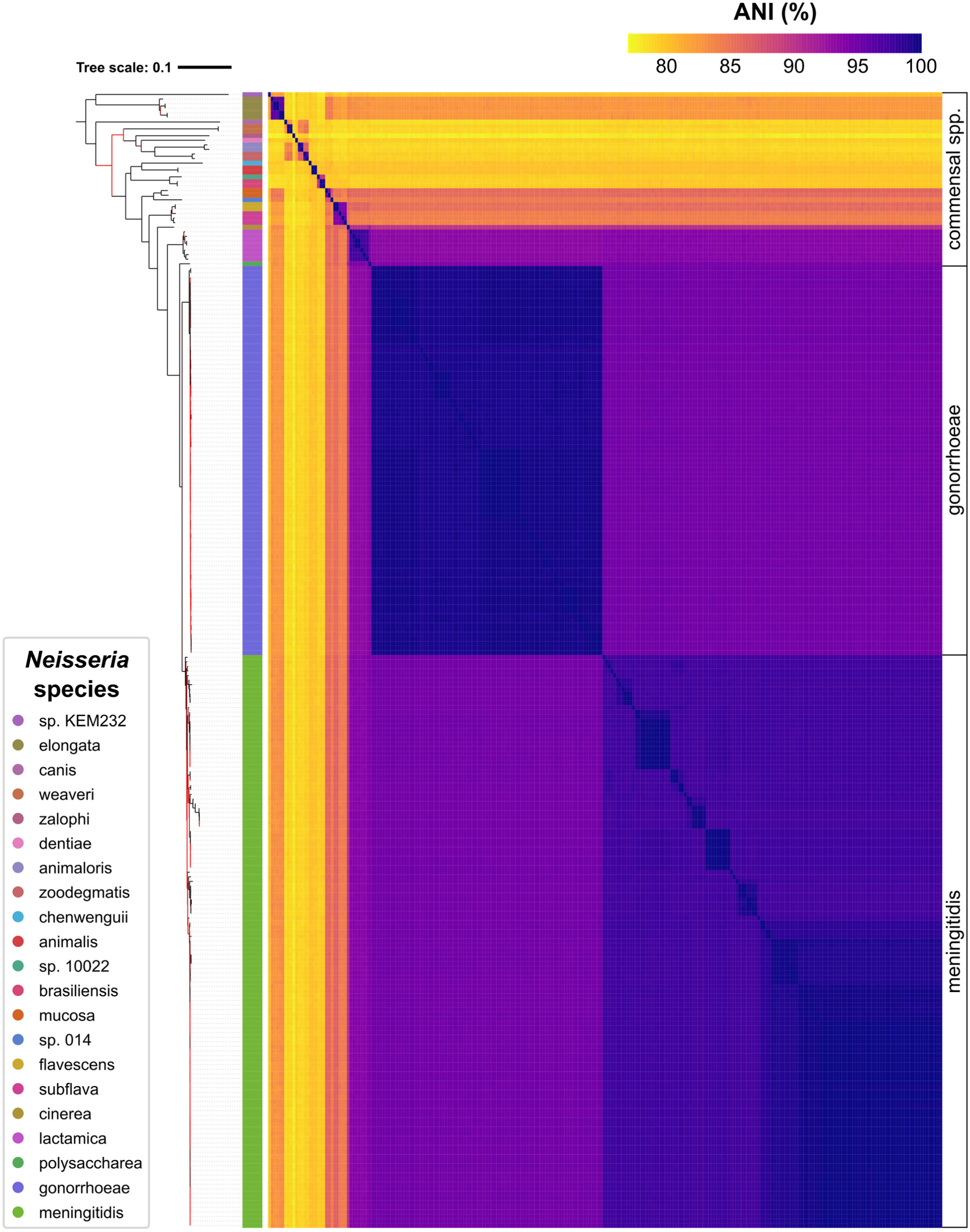
Relationships between bacterial genomes included in this study. Maximum-likelihood ribosomal MLST (rMLST) tree of the smaller set of high-quality *Neisseria* genomes (described in Table S1A) and a heatmap of pairwise average nucleotide identity (ANI) values between each genome. Ribosomal gene sequences of each genome were identified and concatenated using PubMLST and then used to create a phylogenetic tree using RAxML. The tree was visualized with iTOL and rooted using midpoint rooting. Each colored bar represents a different *Neisseria* species as defined in the key; the order of species in the tree is the same as the order shown in the key. Bootstrap support is indicated by the color of each branch, where red indicates low support. The tree file is provided in Data Set S1, Tab 1. ANI values were calculated using FastANI and represented as a color gradient as indicated in the key.

To increase the likelihood of identifying novel phages, we selected 3 tools that use different approaches to predict prophages within *Neisseria* genomes. PhiSpy uses machine learning to search for characteristics that are unique to prophages (47), VirSorter2 combines alignment and machine learning-based approaches to identify microbial viruses (48, 49), and Seeker uses deep learning to detect phages without relying on any sequence features (50).

We assessed whether these tools could identify 9 known *Neisseria* prophages (**Table S1B**) in *N. gonorrhoeae* FA 1090 (24, 26, 28) and 4 in *N. meningitidis* Z2491 (24, 25, 29). Combined, these three tools predicted 6/9 known, intact prophages in FA 1090 (**Fig. S1A**), and 2/4 in Z2491 (**Fig. S1B**). None of these tools correctly identified known *Neisseria* filamentous prophages; they were either missed entirely or combined with an adjacent dsDNA prophage into a single prediction (NgoΦ2 with NgoΦ6; NgoΦ3 with NgoΦ9) (**Fig. S1A**). These results are consistent with previous observations that several tools (including VirSorter2) have difficulty predicting *Neisseria* prophages (51).

In total, we obtained 2050 predictions (**Table S2A**), which were reduced to 1302 unique predictions after dereplication at 95% length aligned (see Methods and **Table S2B**). No phages identified in different bacterial species were found to be similar at ≥95% length aligned (**Table S2B**). The distribution of lengths of dereplicated prophages predicted by each tool is shown in **Fig. S2**.

Subsequently, we present analyses on predictions made by all 3 tools. For the sake of space and clarity, analyses of PhiSpy predictions are presented in the main text, while analyses of VirSorter2 and Seeker predictions are included in the Supplemental Material. We focus on a single tool to avoid the issue of reconciling overlapping predictions between tools and selected PhiSpy because it more accurately predicted the boundaries of known *Neisseria* prophages (**Fig. S1**).

### Few predictions are similar to *Neisseria* plasmids and the Gonococcal Genetic Island

In this study, we used prediction tools that search for viruses. However, because VirSorter2 has been reported to have difficulty distinguishing plasmids from viral sequences (49, 52, 53), we wanted to address the possibility that predictions from any of the tools may resemble other types of *Neisseria* MGEs.

Specifically, we compared our predictions to known *Neisseria* plasmids and the Gonococcal Genetic Island (GGI). To perform this analysis, we performed hierarchical clustering based on percent length aligned of dereplicated predictions and the nucleotide sequences of plasmids and GGI obtained from GenBank (44, 45) (**Table S1C**).

Only 14 unique predictions cluster with *Neisseria* plasmids and the GGI based on nucleotide sequence (**Fig. S3, Data Set S1**). Of these 14 predictions, 2 of them (both predicted by VirSorter2) cluster with known *Neisseria* plasmids (**Fig. S3A**), and 12 predictions (all predicted by Seeker) cluster with the GGI (**Fig. S3B**).

Because our study is focused on phages, we excluded these 14 predictions from our subsequent analyses. The results above indicate that the majority of predictions in this study are dissimilar from these other types of known *Neisseria* MGEs.

### Comparing predictions to known phages using gene-sharing networks

Classifying phages is challenging due to their high genomic diversity, extensive mosaicism, and lack of universally shared genes (54–57). Therefore, gene-sharing networks are commonly used to compare novel phages to previously identified phages (35, 36, 57–61). Here, we used vConTACT v.2.0 (60, 62) to assess whether the prophages we predicted are similar to known *Neisseria* phages (**Table S1B**) or phages that infect other bacterial taxa (i.e., other reference viruses; see Methods for details).

vConTACT generates a similarity score between each pair of viruses based on the protein clusters they share. If two viruses are significantly similar to one another (i.e., the pair has a score ≥1), then they are connected by an edge. Groups of viruses that are highly similar are placed within the same subcluster and are likely members of the same viral genus (60). We used vConTACT to separately analyze dereplicated predictions from each tool, resulting in 3 distinct networks (**Fig. 2, Fig. S4A and S4B**).

**Fig. 2.**
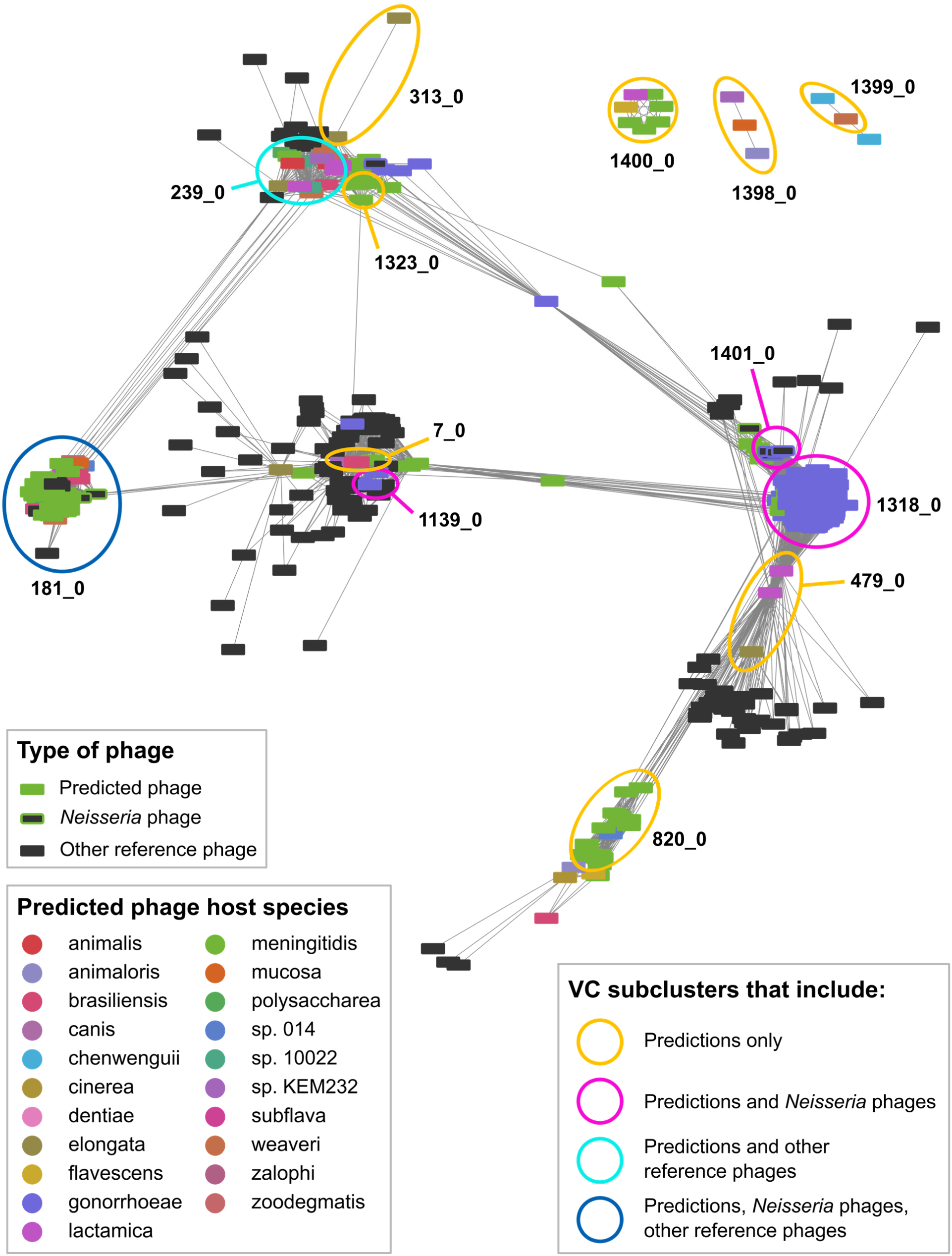
vConTACT clustering of PhiSpy predictions, *Neisseria* phages, and phages that infect other bacterial taxa. vConTACT v.2.0-generated network of dereplicated PhiSpy predictions and reference viruses visualized with Cytoscape using an edge-weighted spring-embedded algorithm. Nodes represent predicted prophages (color corresponding to the *Neisseria* species in which prophages were identified), *Neisseria* phages (dark grey outlined in the color corresponding to the bacterial host species), or other reference viruses (dark grey without outline). Edges represent the vConTACT-generated similarity score between each pair of viruses (only similarity scores ≥1 are included in the network). Highly similar viruses are positioned close together. Only reference viruses that are connected to ≥1 predicted prophage are included in the network. Information about vConTACT subclusters is included in Data Set S1, Tab 3, and similarity scores (edge weights) in Tab 4.

First, we examined whether PhiSpy predictions are significantly similar to known *Neisseria* phages (i.e., connected by an edge in the network). While 83% of PhiSpy predictions (229/277) are *connected to* known *Neisseria* phages, only 52% of PhiSpy predictions (144/277) *cluster with Neisseria* phages (**Table S2C**). These 144 predictions belong to the following 4 subclusters: 181_0 (dark blue circle), 1139_0, 1401_0, 1318_0 (pink circles, **Fig. 2**). Thus, only half of PhiSpy predictions are likely members of the same viral genus as known *Neisseria* phages.

Next, we compared PhiSpy predictions to viruses that infect bacterial taxa other than *Neisseria* (i.e., other reference viruses). We found that 86% of PhiSpy predictions (239/277) are significantly *connected to* reference viruses (**Table S2C**). However, only 15% of PhiSpy predictions (42/277) *cluster with* other reference viruses (**Table S2C**); these predictions belong to either subcluster 181_0 (dark blue circle) or 239_0 (light blue circle, **Fig. 2**). Below, we explore these 2 subclusters that contain both PhiSpy predictions and other reference viruses.

Subcluster 181_0 includes Mu-like phages that infect *N. meningitidis* (Pnm1-2, MuMenB), *Mannhemia haemolytica* (3927AP2), and *Haemophilus parasuis* (SuMu, shown in bold, **Fig. 3A**). Previously, Pnm1-2 and MuMenB were found to resemble a Mu-like phage that infects *H. influenzae* (21). There is a high degree of synteny between members of 181_0, and the proteins shared between known and predicted *Neisseria* prophages in this subcluster have >50% sequence identity (**Fig. 3A**).

**Fig. 3.**
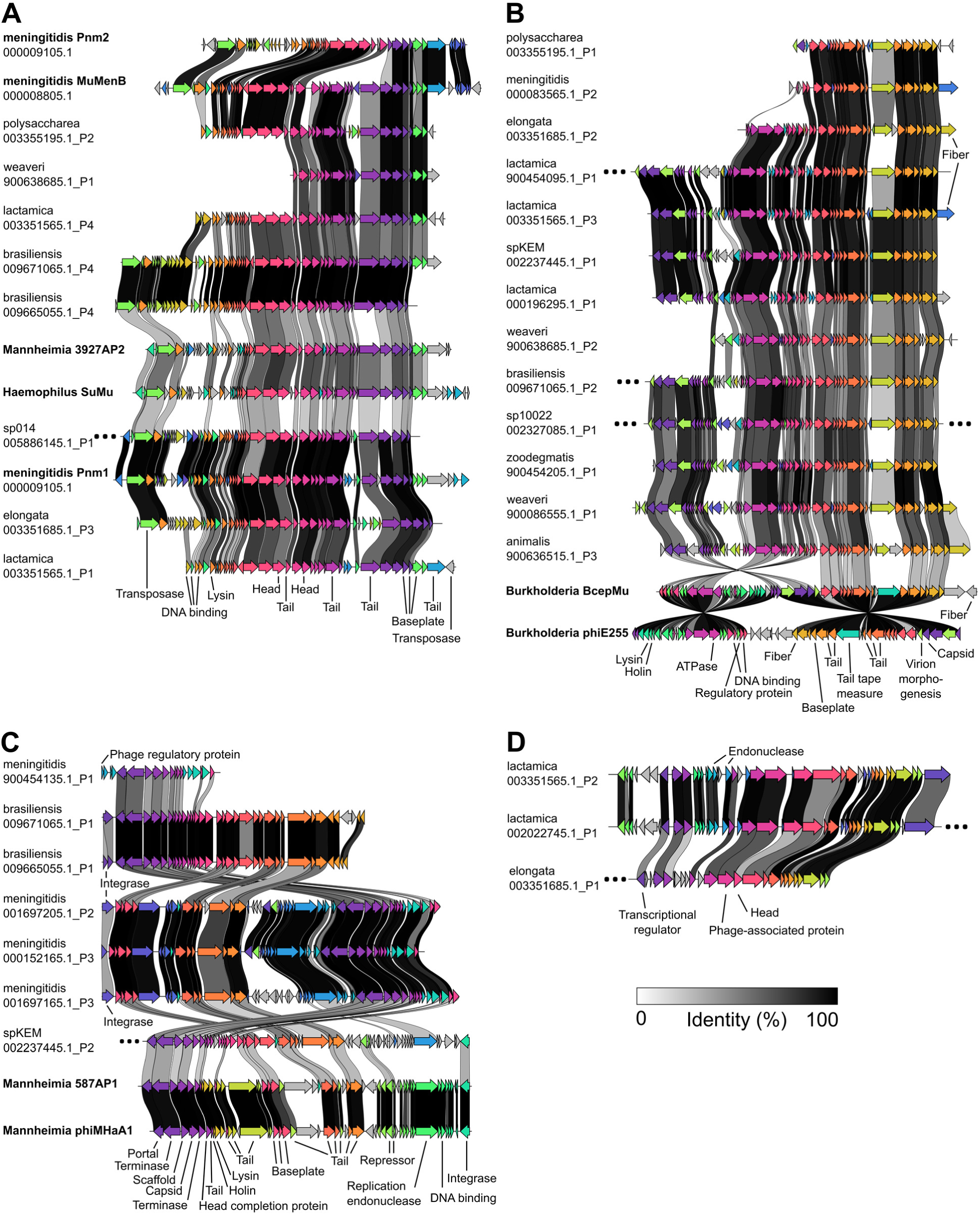
Shared genes between predictions and reference phages. (A-D) Clinker-generated visualizations showing genes that are shared between members of the following vConTACT subclusters: 181_0 (A), 239_0 (B), 7_0 (C), and 479_0 (D). The location of each subcluster within the vConTACT-generated network is shown in Fig. 2. Reference phages are indicated in bold. Arrows with the same color indicate genes that are similar between phages; connections between arrows indicate amino acid sequence identity as described in the key. Grey arrows indicate genes that are not shared between phages; ellipses indicate that >2 unshared genes are present at either end of a phage genome. Shared genes are annotated with predicted functions of encoded proteins (except for hypothetical proteins). In Panel A, several late phage genes are present/absent between *Burkholderia* phages and *Neisseria* predictions (their order in phiE255 from left to right: lysin, holin, tail tape measure, and fiber proteins). Panel C also includes two reference viruses that are not part of the subcluster: *Mannheimia* phages 587AP1 and phiMHaA1. Information about vConTACT subclusters is included in Data Set S1, Tab 3.

Similarly, subcluster 239_0 contains two other Mu-like phages that infect *Burkholderia cenocepacia* (BcepMu) and *B. thailandensis* (phiE255, **Fig. 3B**). Except for several late phage genes, most predicted proteins are shared between the *Burkholderia* phages and *Neisseria* predictions (at ∼30-50% sequence identity, **Fig. 3B**). Together, the results above suggest that several *Neisseria* species may be infected by phages that are similar to those that infect *Haemophilus, Mannheimia*, and *Burkholderia* — microbes that *Neisseria* species may encounter within the respiratory tracts of humans and animals.

We also explored subcluster 7_0 (orange circle, **Fig. 2**). Although this subcluster does not include any reference viruses, its members share many genes with reference viruses (many surrounding dark grey nodes, **Fig. 2**). In particular, a *Neisseria* sp. KEM 232 prediction that is part of 7_0 shares 48% of predicted proteins (29/60) with other members of 7_0 (**Fig. 3C**), and also shares 30% of proteins (18/60) with two *Mannheimia* P2-like phages that do not belong to this subcluster (587AP1 and phiMHaA1, **Fig. 3C**).

While the majority of PhiSpy predictions do not cluster with known phages (**Table S2C**), many predictions were found to have a low degree of similarity to different reference phages (as indicated by the low similarity scores between predictions and reference viruses, **Fig. S5**). For example, the *N. lactamica* and *N. elongata* predictions belonging to 479_0 each have a low degree of similarity to ∼20-50 different reference viruses (**Fig. 3D**). Therefore, these findings suggest that several *Neisseria* prophages are distantly related to multiple viruses that infect other bacterial taxa.

Finally, we found differences in how similar predictions from each tool are to known viruses. Specifically, 1) a smaller proportion of Seeker predictions are *connected to* reference phages compared to PhiSpy and VirSorter2 predictions (**Table S2C**); 2) the *degree to which* Seeker predictions are similar to reference viruses is significantly lower compared to the other tools (**Fig. S5**); and 3) zero Seeker predictions *cluster with* reference viruses (**Table S2C**). Thus, Seeker predictions may represent novel phages, other MGEs, or alternatively, regions of the chromosome that were incorrectly called.

### Highly similar predicted prophages are found in distantly related *Neisseria* species

Next, we explored whether different *Neisseria* species may be infected by highly similar phages by examining whether any vConTACT clusters include PhiSpy predictions found in genomes of different *Neisseria* species (**Fig. 4A**).

**Fig. 4.**
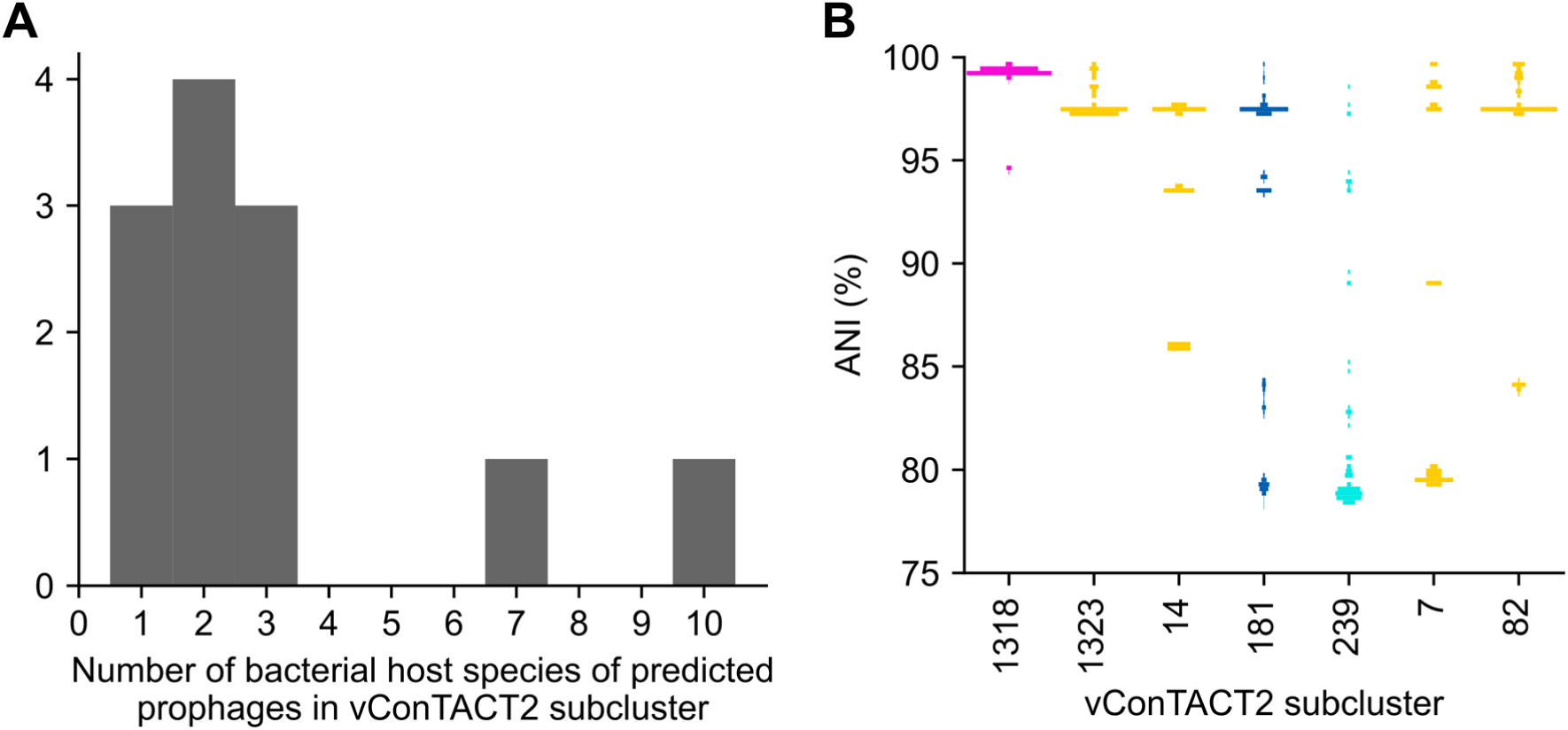
Several clusters include PhiSpy predictions identified in different host bacterial species. (A) The number of different host bacterial species of PhiSpy predictions within each vConTACT v2.0 subcluster. (B) The percent average nucleotide identity (ANI) between the bacterial genomes in which PhiSpy predictions were identified. Only vConTACT subclusters that include 5 or more predicted prophages are shown (“_0” was omitted from the end of subcluster names). The distribution of ANI values is represented as a histogram where the width of bars at a given ANI corresponds to the proportion of genome pairs with that ANI. The color of each subcluster indicates the types of predictions that belong to that subcluster (orange: predictions only; pink: predictions and *Neisseria* phages; light blue: predictions and other reference phages; dark blue: predictions, *Neisseria* phages, and other reference phages). Information about vConTACT subclusters is included in Data Set S1, Tab 3.

We found that out of the 12 subclusters that include PhiSpy predictions, 9 contain predictions identified in different *Neisseria* species (**Fig. 4A**). Strikingly, subclusters 181_0 and 239_0 (**Fig. 3A and B**) include predictions found in 7 and 10 different *Neisseria* species, respectively (**Fig. 4A**).

Afterwards, we investigated whether the bacterial species (in which these predictions were identified) are closely or distantly related to each other. For every subcluster that includes ≥5 predictions, we calculated the pairwise average nucleotide identity (ANI) between each bacterial genome in which the predicted prophages were identified.

Every subcluster we examined includes multiple phages found in the same species, as shown by ANI >95% (**Fig. 4B**). Also, 4 clusters include predictions found in closely related species (ANI 90-95%).

Finally, 5 clusters include predictions found in more distantly related species (ANI <90%, **Fig. 4B**). Strikingly, host bacterial genomes of predictions in 3 of these clusters (181_0, 239_0, 7_0, **Fig. 3A-C**) have ANI ∼80% (**Fig. 4B**). Therefore, these results suggest that even distantly related *Neisseria* species may be infected by closely related phages.

### Identification of CRISPR arrays and matches within *Neisseria* genomes

Here, we surveyed a larger set of 2619 *Neisseria* genomes (see Methods, **Table S1D**) for the presence of CRISPR arrays (see **Data Set S1**).

Consistent with previous findings (40–42), we identified Type II-C CRISPR arrays in 45% of *N. meningitidis* genomes (862/1894), and no CRISPR arrays in *N. gonorrhoeae* genomes (0/630). Also, we identified CRISPR arrays in ≥1 genome of every commensal species included in this study. Repeat sequences in arrays of commensal species are associated with 7 different CRISPR subtypes (I-A, I-C, I-F, II-C, III-A, III-B, III-D). In total, we found 3676 unique CRISPR spacers.

Next, we used BLASTn to search for matches between CRISPR spacers and sequences within *Neisseria* genomes. We only kept matches that had 100% identity over the entire length of the spacer (i.e., 0 mismatches), and we looked for both intra- and inter-species matches. We found that 22% of spacers (800/3676) target sequences within the smaller set of high-quality *Neisseria* genomes. Out of these targeting spacers, 67% (539/800) match known or predicted prophages.

Previously, Zhang et al. identified 5 self-targeting spacers in 6 *N. meningitidis* genomes (41). Here, we found that 52% of CRISPR-positive *N. meningitidis* genomes (23/44) encode self-targeting spacers.

### Examining the locations of CRISPR matches within *Neisseria* genomes

We next examined the genomic locations of spacer matches. In addition to providing defense against MGEs, the Type II-C CRISPR system of *N. meningitidis* has been proposed to play a role in limiting natural transformation (41, 42). If CRISPR systems limit transformation, we would expect to see targeting evenly distributed along the length of the bacterial chromosome with no obvious enrichment of targeting in any location. If, however, prophages are targeted by CRISPR immunity, we would expect that matches would be enriched within prophages.

**Fig. 5** shows the genomic locations of spacer matches within 2 genomes that encode CRISPR arrays (*N. meningitidis*, sp. 10022) and 2 that do not (*N. gonorrhoeae, N. weaveri*). In *N. gonorrhoeae* and *N. meningitidis*, we observe a low level of targeting across the length of the genome (**Fig. 5A**), consistent with a role for CRISPR systems in restricting transformation. There are also regions of high targeting: in *N. gonorrhoeae*, peaks correspond to the location of several known prophages, whereas peaks in the *N. meningitidis* genome may correspond to as of yet unidentified MGEs (**Fig. 5A**).

**Fig. 5.**
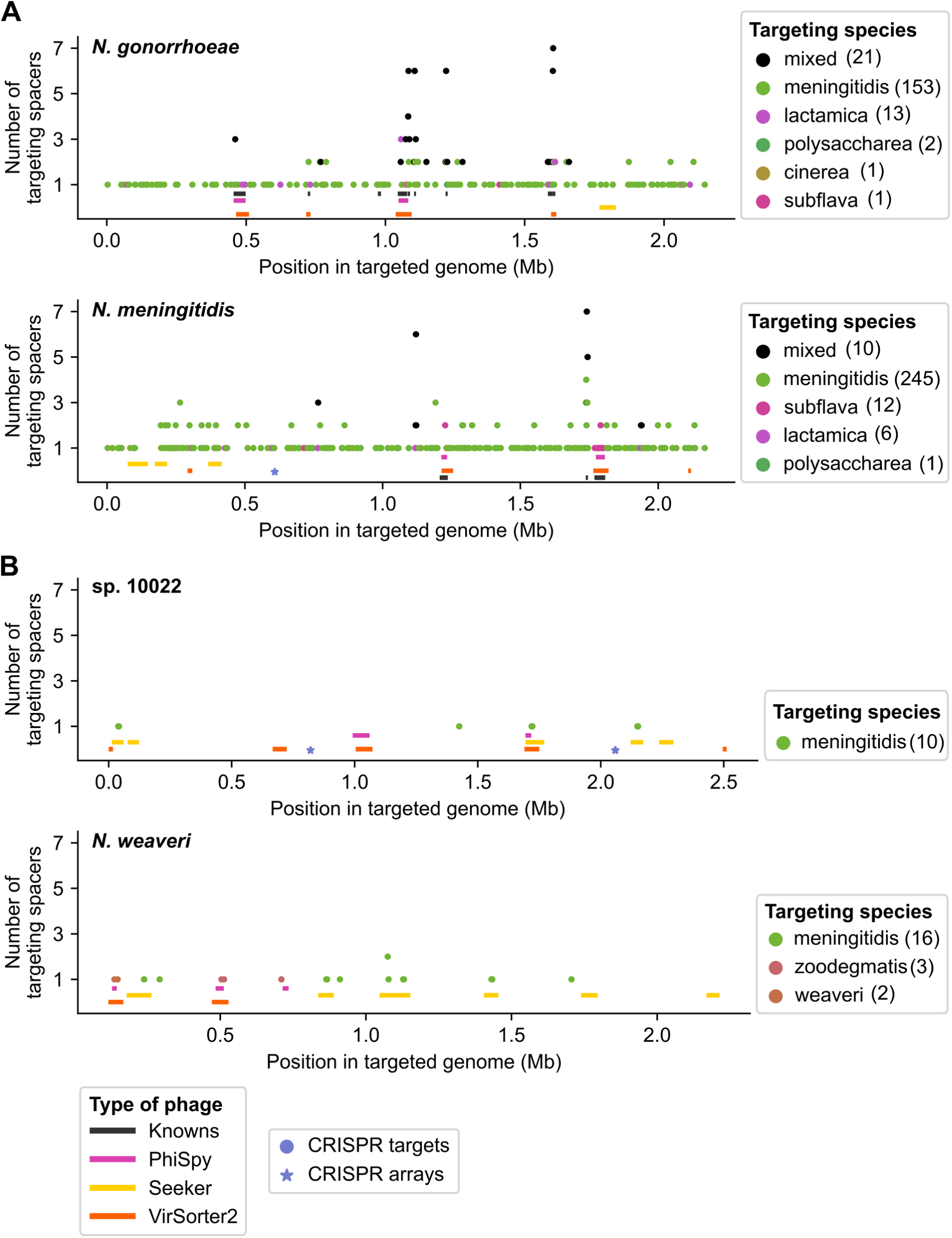
Locations of matches between CRISPR spacers and *Neisseria* genomes. (A and B) Genomic locations of matches between *Neisseria* CRISPR spacers and bacterial genomes. Each plot shows a genome of a different species: *N. gonorrhoeae* (GCA_000006845.1) and *N. meningitidis* (GCA_000009105.1) (A), and sp. 10022 (GCA_002327085.1) and *N. weaveri* (GCA_900638685.1) (B). Each data point (circle) represents the number of spacers that match each position (1-kb bin) in the bacterial genome. The color of the circle corresponds to the species encoding the spacer (as indicated in the key, with mixed indicating >1 unique species); the number in parentheses indicates the number of spacers encoded by each species that match the genome (at 100% identity over the entire spacer length). CRISPR arrays are denoted by stars, and rectangles represent prophages (as indicated in the key). Information about CRISPR targeting of each genome is provided in Data Set S1, Tab 10; locations of CRISPR arrays are in Tab 7; and locations of known and predicted prophages are in Tables S1B and S2A, respectively.

Matches within sp. 10022 and *N. weaveri* genomes mostly correspond to predicted prophages (**Fig. 5B**). Overall, many fewer spacers appear to target sp. 10022 and *N. weaveri* genomes, which is due at least in part to the few available genomes of commensal species, leading to a small pool of targeting spacers. Thus, our ability to make comparisons of targeting between *N. meningitidis* and commensals is limited.

Finally, we examined which species encode the targeting spacers (color of each circle, **Fig. 5**). Previously, *N. meningitidis* spacers were reported to match protospacers in *N. gonorrhoeae* genomes (41); here, we observe that *N. meningitidis* is largely responsible for the low-level targeting of the *N. gonorrhoeae* and *N. meningitidis* genomes in **Fig. 5A**. In contrast, prophages in these 4 genomes are matched by spacers from *N. meningitidis* or other species (**Fig. 5**). In subsequent analyses, we quantify CRISPR targeting of each prophage and bacterial genome and further investigate interspecies targeting.

### Comparing CRISPR targeting of each predicted prophage to backbone targeting

Above, we observed CRISPR targeting along the entire length of the chromosome in *N. gonorrhoeae* and *N. meningitidis*. To distinguish whether predicted prophages are preferentially targeted, it is necessary to compare the level of targeting of predicted prophages to the background level across the rest of the genome. Therefore, we quantified the density of CRISPR targeting of every predicted prophage and the rest of the bacterial genome in which it was identified (i.e., the backbone).

We defined prophage targeting density as the number of CRISPR matches within the prophage divided by the prophage length. The backbone targeting density was obtained by dividing backbone targeting (the number of CRISPR matches within a bacterial genome excluding targets in all known or predicted prophages and CRISPR arrays) by the length of the backbone (length of the entire bacterial genome minus the combined lengths of the prophages identified in that genome).

First, we compared the targeting density of each prophage to the targeting density of the backbone genome (**Fig. 6A**). We then compared ratios of prophage/backbone targeting between different *Neisseria* species (**Fig. 6B**). Although many *N. gonorrhoeae* and *N. meningitidis* prophages have high targeting densities (**Fig. 6A**), the high degree of backbone targeting of these genomes results in mostly low targeting ratios (**Fig. 6B**). For example, even though all 13 known *Neisseria* prophages are matched by spacers, only 5 of them (MDAΦ and NgoΦ6-9) are significantly more highly targeted compared to the backbone (**Data Set S1**).

**Fig. 6.**
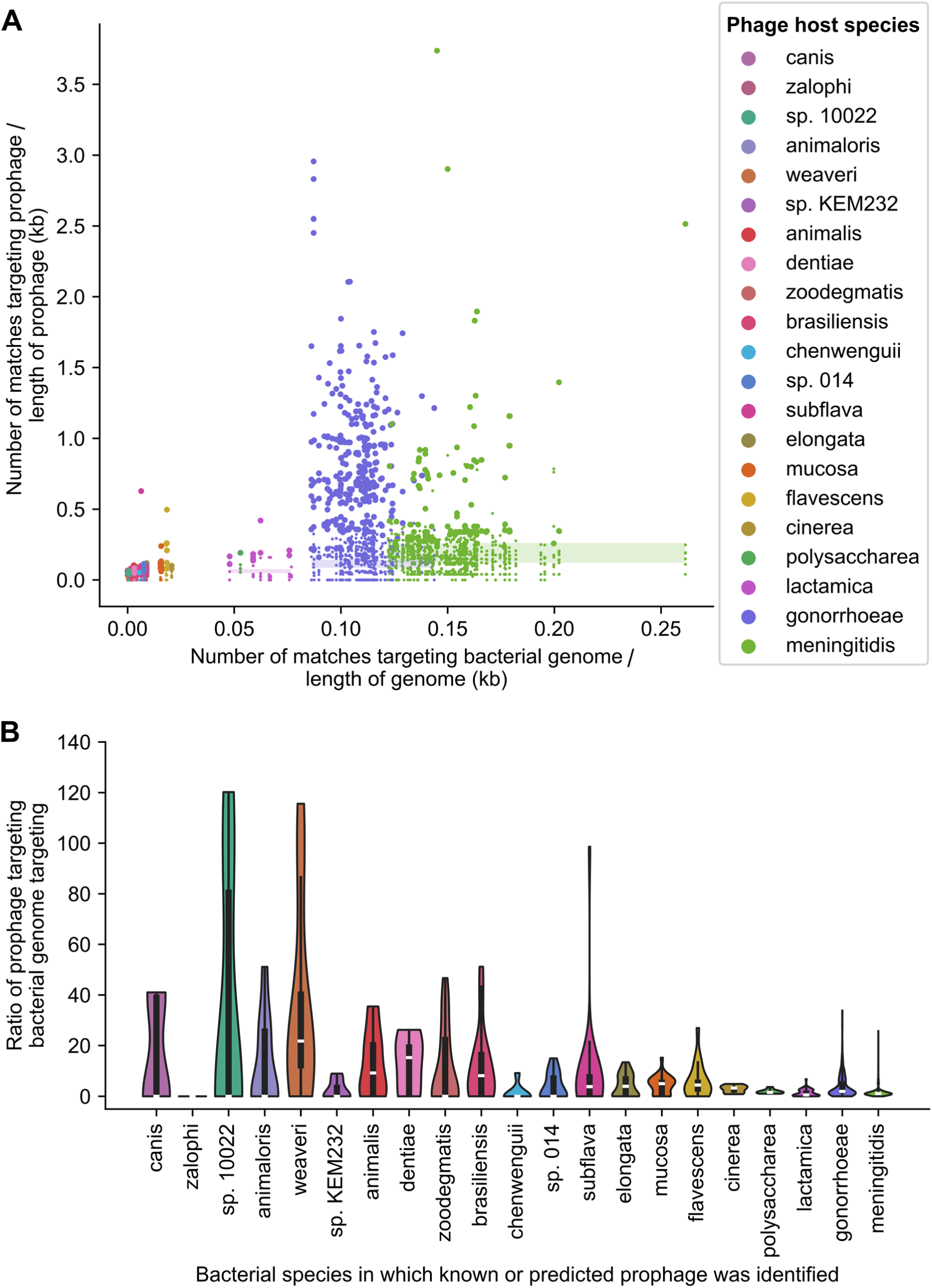
Comparing CRISPR targeting density of prophages to targeting of bacterial genome backbones. (A) Matches were identified between CRISPR spacers and bacterial genomes (at 100% identity over the entire spacer length). For each prophage, the density of matches within the prophage was compared to the backbone genome in which the prophage was identified (targeting densities are provided in Data Set S1, Tab 10). Y coordinates represent the number of matches within each prophage divided by the length of the prophage. X coordinates represent the number of matches within the bacterial genome in which the prophage was identified (excluding spacers targeting all prophages that were predicted in that genome) divided by the length of the bacterial genome (minus the length of all predicted prophages within that genome). Each data point represents a known *Neisseria* prophage or a predicted prophage. The color of the circle corresponds to the species in which the prophage was identified. Larger circles indicate prophages that have significantly higher targeting densities compared to the backbone (as described in the Methods and listed in Data Set S1, Tab 11). To highlight which prophages are targeted more highly than the backbone sequence, the range of targeting densities for genomes of each species is shown as a translucent box; the color of the box corresponds to the species. (B) Distribution of prophage/backbone CRISPR targeting ratios grouped according to the species in which the prophage was identified. Only dereplicated predicted prophages are included. CRISPR targeting ratios are provided in Data Set S1, Tab 10.

In multiple commensal species, the ratio of prophage/backbone targeting is very high (**Fig. 6B**), in many cases due to little or no backbone targeting (**Fig. 6A**). Low levels of backbone targeting could be due to several, non-mutually exclusive reasons: 1) the small number of commensal spacers sampled in this study resulting in lower apparent targeting; 2) infrequent encounters between certain species (e.g., *N. weaveri* is an opportunistic pathogen rather than a resident of the human mucosa) (63); or 3) species-specific barriers to transformation, including differences in DNA uptake sequences (64) and in whether CRISPR systems target chromosomal DNA.

Overall, 20% of dereplicated prophages predicted in this study (259/1306) have a significantly higher targeting density compared to the backbone (**Table S2C, Data Set S1**). Furthermore, the majority of significantly targeted predictions (74%; 191/259) do not cluster with known *Neisseria* phages, plasmids, or the GGI (**Table S2C**); these predictions belong to 30 different vConTACT subclusters (**Data Set S1**), including 3 of the PhiSpy subclusters highlighted above (239_0, 7_0, 479_0, **Fig. 3**). Therefore, these 191 predictions represent likely candidates for novel *Neisseria* phages.

### Interspecies CRISPR targeting is widespread among *Neisseria* species

We observed a high degree of interspecies targeting in our dataset. Out of 539 spacers that target known or predicted prophages, 53% (288/539) are involved in interspecies targeting of prophages. Furthermore, 35% (186/539) *only* target prophages found in other species.

We further examined interspecies CRISPR targeting using a network representation of targeting relationships between *Neisseria* species (**Fig. 7**). Specifically, the network includes targeting of backbone sequences, known *Neisseria* prophages, and predicted prophages that are more highly targeted compared to the backbone and that do not cluster with *Neisseria* plasmids.

**Fig. 7.**
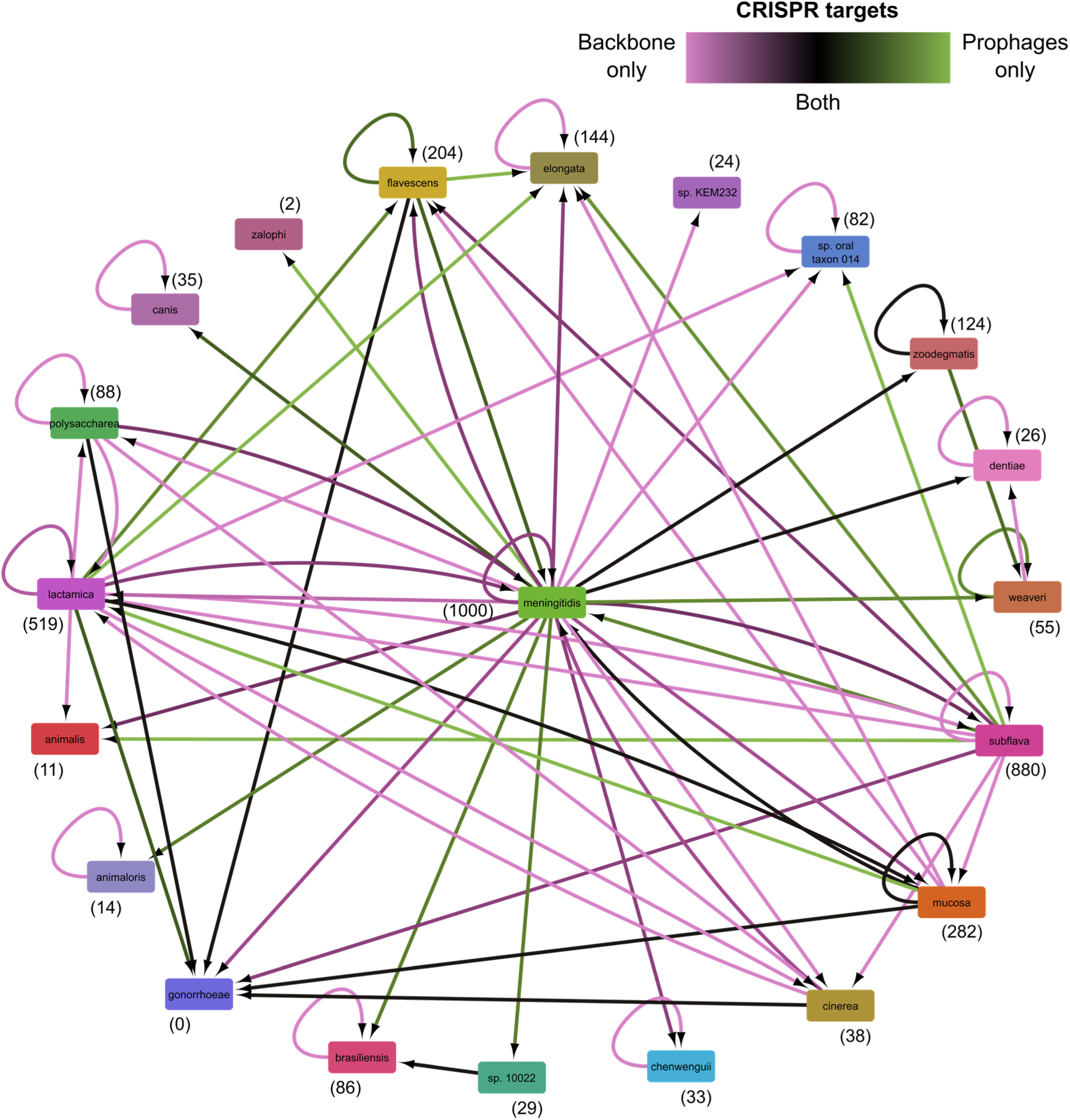
Interspecies CRISPR targeting of *Neisseria* prophages and bacterial genome backbones. Network representing intra- and inter-species CRISPR targeting of prophages and bacterial backbone sequences visualized using Cytoscape. Each node represents a bacterial species, and the adjacent number in parentheses indicates the total number of spacers encoded by that species. Nodes are connected by an edge if CRISPR spacers encoded by one species target another species. The direction of CRISPR targeting is indicated using an arrow that points to the species being targeted. Edge color indicates the relative number of spacers targeting prophages compared to the targeting of the backbone (i.e., bacterial genome excluding CRISPR arrays or sequence contained within any known or predicted prophages — not only prophages that are significantly targeted). Included in this analysis are backbone sequences, known *Neisseria* prophages, and predicted prophages that are targeted significantly more than the backbone and do not cluster with *Neisseria* plasmids. Information about CRISPR spacers and targeting is provided in Data Set S1, Tabs 9-10.

The network is highly interconnected: all 21 *Neisseria* species included in this study are connected to ≥1 other species in the network, and 16 species are connected to ≥2 others (**Fig. 7**). Interestingly, *N. meningitidis* spacers match prophage and backbone sequences of 17 and 20 different species, respectively. Additionally, *N. gonorrhoeae* and *N. meningitidis* are each targeted by 7 *Neisseria* species.

Moreover, there are differences in the type of sequences targeted (edge color, **Fig. 7**). *N. meningitidis* spacers seem to predominantly target the backbone genome sequences of *N. meningitidis* and several other species (many pink arrows from *N. meningitidis*). In contrast, *N. subflava* and *N. lactamica* spacers primarily target prophages of other species (green arrows).

While these results suggest that interspecies targeting of *Neisseria* sequences is widespread, an alternative explanation is that spacers were exchanged between species. However, out of 3676 total spacers, only 2 identical spacers were present in genomes from different species (**Data Set S1**). Taken together, the findings above indicate that interspecies CRISPR targeting is common between *Neisseria* species.

Finally, we investigated whether *Neisseria* prophages may be targeted by other bacterial taxa using the tool CRISPRopenDB and its database of 11 million spacers (65). Four predictions identified in *N. animalis* are matched by a single spacer from *Eikenella corrodens* (another member of the Neisseriaceae), while a *N. elongata* prediction is matched by one *Aggregatibacter aphrophilus* spacer (**Data Set S1**).

### Many known and predicted *Neisseria* prophages have additional inferred host species

Elucidating the host range of phages is critical for understanding how they influence their microbial hosts, including their role in mobilizing DNA (12). CRISPR targeting data are frequently used to predict the bacterial hosts of phages (35–39); therefore, we took advantage of the extensive interspecies CRISPR targeting observed above (**Fig. 7**) to infer the hosts of known *Neisseria* prophages and predictions from all 3 tools.

To increase the likelihood of examining true prophages (instead of chromosomal sequences), we only included dereplicated predictions that have significantly higher targeting densities compared to the backbone (i.e., significantly targeted predictions) identified above (**Fig. 6**).

We found that 98% of significantly targeted predicted prophages are likely shared with ≥1 other species (254/259) (**Fig. 8A, Table S2D**). Additionally, 100% of significantly targeted *N. gonorrhoeae* predictions (145/145) and 93% of significantly targeted *N. meningitidis* predictions (26/28) have ≥2 inferred additional host species. Furthermore, 50% (72/145) of predicted *N. gonorrhoeae* prophages have ≥4 additional host species (**Fig. 8A**).

**Fig. 8.**
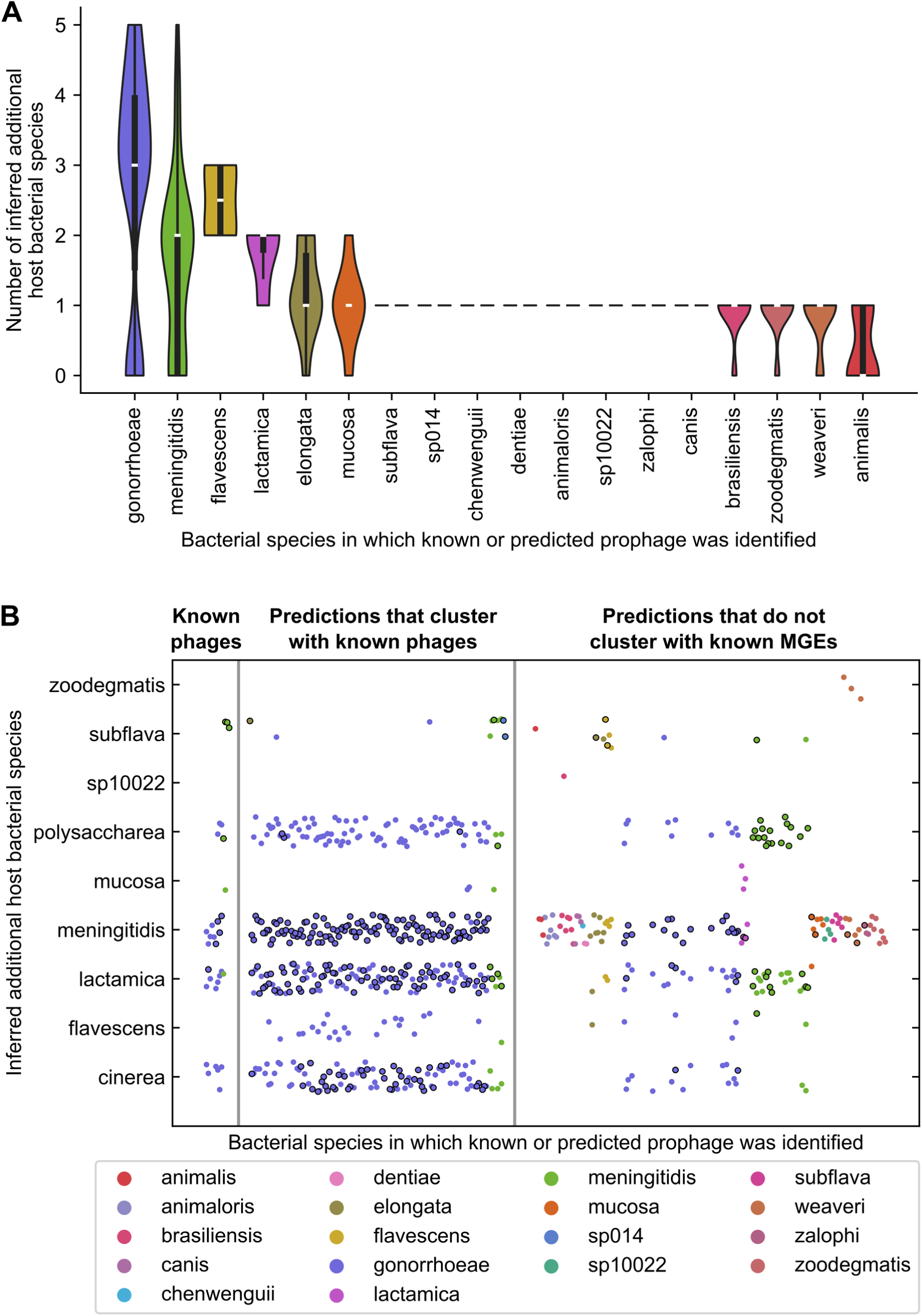
Inference of additional host species of *Neisseria* phages using interspecies CRISPR targeting. (A) Distribution of the number of inferred additional host species of known *Neisseria* phages and dereplicated phages predicted by PhiSpy, VirSorter2, and Seeker (only those that have a significantly higher CRISPR targeting density compared to the bacterial genome backbone, Data Set S1, Tab 11). Interspecies matches between CRISPR spacers and prophages were used to infer the phage hosts. The data are grouped according to the species in which the prediction was identified. (B) Inferred sharing of prophages between the indicated *Neisseria* species. Each circle represents a known or predicted prophage, and the color of the circle corresponds to the species in which the prophage was identified (as indicated in the key). Circles with a black outline indicate that >1 spacer encoded by that species matches that prophage, while circles without an outline represent a single spacer match. Only prophages that have ≥1 inferred additional host species are shown. Predicted prophages are divided into 2 categories: those that cluster with known *Neisseria* prophages and those that do not cluster with known *Neisseria* MGEs (phages, plasmids, or the Gonococcal Genetic Island). The species names along the Y axis are the additional bacterial host species inferred using interspecies targeting data.

Interspecies targeting data suggest that prophages are likely shared between a variety of *Neisseria* species (**Fig. 8B**). *N. gonorrhoeae* prophages (indicated by the many purple dots in multiple rows, **Fig. 8B**) are shared primarily with 3 closely related species, *N. meningitidis, N. polysaccharea*, and *N. lactamica* (median ANI between each one and *N. gonorrhoeae*: 93-95%, **Fig. 1**), and also with *N. cinerea*, which is less closely related to *N. gonorrhoeae* (median ANI: 90%).

Furthermore, *N. meningitidis* is predicted to share phages with a wide range of species, especially predictions that do not cluster with known *Neisseria* MGEs (many differently colored dots in the row corresponding to *N. meningitidis*, **Fig. 8B**). Interestingly, 7 of these species (*N. animaloris, N. canis, N. dentiae*, sp. 10022, *N. weaveri, N. zalophi, N. zoodegmatis*, **Fig. 8B**) are distantly related to *N. meningitidis* (median ANI: <80%, **Fig. 1**).

Finally, we compared the number of inferred additional host species of PhiSpy predictions between vConTACT subclusters. Significantly targeted members of 1318_0 (which includes the known prophages NgoΦ1 and NgoΦ2) have 3-5 additional host species, while members of other subclusters have 0-3 (**Fig. S6**). Together, our findings suggest that diverse *Neisseria* species may be infected by the same phages.

## DISCUSSION

In this study, we sought to broaden the diversity of phages known to infect *Neisseria* species. We used 3 different virus prediction tools to scan 248 genomes of commensal and pathogenic *Neisseria* species for prophages. Clustering approaches revealed that many of these predictions are dissimilar from known *Neisseria* MGEs (prophages, plasmids, or the GGI) and phages described to infect other taxa. Therefore, we may have uncovered novel *Neisseria* prophage diversity.

We also identified prophages in several commensal *Neisseria* species that are highly similar to the *N. meningitidis* prophages Pnm1-2 and MuMenB, as well as Mu-like phages that infect *Haemophilus parasuis* and *Mannheimia haemolytica*, two Gammaproteobacteria. Although Pnm2 and MuMenB are defective, they may retain the ability to contribute genes to other co-infecting phages (55). Interestingly, several predicted *Neisseria* prophages are highly similar to Mu-like phages that infect other Betaproteobacteria: *Burkholderia cepacia* and *B. thailandensis*.

Commensal *Neisseria* species frequently colonize the upper respiratory tracts of humans and animals. Thus, these species may encounter *Haemophilus, Mannheimia*, and *Burkholderia* species within these niches and could be exposed to the same or similar phages. Additionally, we found highly similar predicted prophages in different *Neisseria* species, including distantly related species.

CRISPR systems provide immunity against MGEs and can be used as a historical record of encounters between microbes and MGEs. Previously, Zhang et al. identified CRISPR spacers that match the filamentous phage MDAΦ (41), and we found that all 13 known *Neisseria* prophages that we examined are matched by spacers. In addition to defending against MGEs, CRISPR systems of *N. meningitidis* and other microbes may also play a role in restricting the exchange of chromosomal DNA between species (41, 42, 66, 67). Our observations that backbone sequences of many *Neisseria* species are targeted by *N. meningitidis* spacers at a low level are consistent with this model.

To identify predictions that are likely to be MGEs, we compared targeting of each predicted prophage to targeting of the backbone. We found that 20% of dereplicated predicted prophages (259/1306) have a significantly higher targeting density compared to the backbone, and 74% of these (191/259) do not cluster with known *Neisseria* MGEs. Therefore, these 191 predictions are candidates that warrant further study.

Furthermore, we found evidence of widespread interspecies targeting of predicted prophages by *Neisseria* spacers and used those data to infer additional host bacterial species of each prediction. Specifically, we focused on predictions that are significantly more highly targeted than the backbone, which are more likely to be actual prophages. Building upon previous findings (33, 34), our results suggest that multiple known and predicted phages may be able to infect multiple species of *Neisseria*, including species that are distantly related.

This study has several important limitations. Commensal species are underrepresented among the available *Neisseria* genome assemblies, and thus, also in this study. This underrepresentation limits our ability to compare patterns of CRISPR targeting between species. Additionally, our inference of additional host species is limited by whether genomes encode CRISPR spacers (e.g., *N. gonorrhoeae* genomes do not encode CRISPR arrays). Moreover, we only used 3 virus prediction tools and among them only PhiSpy was specifically designed to predict prophages (52).

Prophages are known to influence the fitness and virulence of many bacterial species, including *N. meningitidis* (15–20). Furthermore, considerable evidence suggests that accessory genes are shared extensively between *Neisseria* species and that commensal species are a reservoir of AR and virulence genes (43, 68–72). Therefore, it is critical to understand the diversity and host range of phages, which have the potential to mobilize genes among *Neisseria* species and alter their evolutionary trajectories. Further research on *N. gonorrhoeae* phages is also crucial for developing phage therapy approaches (73).

By combining clustering and CRISPR targeting analyses, we have identified candidate, novel *Neisseria* phages and inferred that several may infect multiple species within this bacterial genus. We hope that our findings may inform future studies seeking to elucidate the impact of viruses on *Neisseria* biology. Finally, we believe that our work may have implications for understanding the interactions occurring among the diverse *Neisseria* species that colonize the oropharynx and the phages that infect them.

## METHODS

### Generation of bacterial genome datasets

A set of high-quality bacterial genome assemblies was selected for prophage prediction. Specifically, *Neisseria* genome assemblies with N50 ≥250 kb and contigs ≤10 were downloaded from GenBank (44, 45). This set of 248 assemblies is referred to as the ‘smaller *Neisseria* genome dataset’ and is described in **Table S1A**. Additionally, for the identification of CRISPR arrays, a second dataset of bacterial genome assemblies was compiled as follows: *Neisseria* genome assemblies with N50 ≥15 kb were downloaded from GenBank (44, 45) and limited to the species represented in the smaller genome dataset. This second set of 2619 assemblies is referred to as the ‘larger *Neisseria* genome dataset’ and is described in **Table S1D**. All assemblies described above were downloaded on 3/30/2020.

### Construction of bacterial phylogenetic tree and heatmap of average nucleotide identity

PubMLST was used to concatenate sequences of the 53 genes encoding ribosomal proteins of each bacterial genome within the smaller *Neisseria* genome dataset (74, 75). Then, the concatenated protein sequences were used to create a maximum-likelihood ribosomal multilocus sequence typing (rMLST) tree using RAxML (with GTRCAT model and 100 bootstrap replicates) (76) and visualized using iTOL v5 (77). The tree was rooted using midpoint rooting. The pairwise average nucleotide identity (ANI) between bacterial genomes in the smaller dataset was calculated using FastANI (78) and visualized as a heatmap using the R package pheatmap (79, 80).

### Prediction of prophages within bacterial genomes

PhiSpy, Virsorter2, and Seeker were used to predict prophages within the smaller set of *Neisseria* genomes (47, 49, 50). PhiSpy was run in strict mode without HMM searches after training using a custom training set. To generate the custom set, we combined the PhiSpy default reference genomes with the *N. gonorrhoeae* genome FA 1090 (GCA_000006845.1) annotated with proteins from the dsDNA tailed phages (NgoΦ1 – 5) and filamentous phages (NgoΦ6 – 9) (26, 28). We did not add any *N. meningitidis* genomes to the training set because *N. meningitidis* MC58 (GCA_000008805.1) and *N. meningitidis* Z2491 (GCA_000009105.1) were already included in the reference dataset. VirSorter2 and Seeker were run using default settings.

### Dereplication of predicted prophages

An all-by-all BLASTn was performed separately with prophages predicted by each tool (81). Predicted prophages were dereplicated at 95% length aligned using a custom script (blast_average_link_hier_clust_output_clusters.py; https://github.com/Alan-Collins/Neisseria-prophage-paper). Information about dereplicated predictions and the predictions used as their representatives is found in **Table S2B**. Known *Neisseria* plasmids were dereplicated using the same method.

### Hierarchical clustering of predicted prophages with *Neisseria* MGEs based on percent length aligned nucleotide sequence

First, an all-by-all BLASTn was performed on dereplicated predicted prophages, dereplicated known *Neisseria* plasmids, and the Gonococcal Genetic Island (81). Next, a distance matrix was created based on the percent length of aligned (PLA) sequence between pairs of MGEs (distance = 1 - PLA). The Python package SciPy v1.6.1 was then used to perform average/UPGMA linkage clustering on the distance matrix (82, 83). Next, a custom script (identify_blast_clusters.py; https://github.com/Alan-Collins/Neisseria-prophage-paper) was used to extract cluster memberships and extract the tree in Newick format, which was visualized using iTOL v5 (77).

### vConTACT v.2.0 clustering of phages based on shared genes

Prodigal was used to predict the protein-coding genes of known *Neisseria* phages and dereplicated predicted prophages (84). Afterwards, Prodigal-generated protein sequences were clustered with reference viral genomes using vConTACT v.2.0 (60, 62). In addition to viruses from the RefSeq database (45, 85), we used 12,892 virus sequences provided by the Millard lab as reference viral sequences (http://millardlab.org/bioinformatics/lab-scripts/supplementing-and-colouring-vcontact2-clusters/). Protein sequence files and mapping files generated on 5/30/2020 were downloaded on 5/20/2021. Clustering with reference sequences was performed separately on prophages predicted by each virus prediction tool. The following vConTACT settings were used: --rel-mode ‘Diamond’, --db ‘ProkaryoticViralRefSeq94-Merged’ --pcs-mode MCL --vcs-mode ClusterONE. Networks were visualized in Cytoscape v3.8.2 using the edge-weighted spring-embedded layout algorithm, which positions highly similar viruses close together (86). Duplicate edges were removed from the network, and reference viruses are only shown if they are connected by an edge to a known or predicted *Neisseria* prophage.

### Analyses of vConTACT viral clusters

We compared the similarity of predicted prophages to reference viruses between the three tools as follows. First, we examined each prediction’s connections to reference viruses and identified the edge with the highest similarity score. Then, we compared the distributions of similarity scores between each tool using the Mann-Whitney U test implemented in the Python package SciPy (82, 83). To compare gene clusters between the members of each cluster, Prokka was used to predict and annotate ORFs within each phage (using Pfam, TIGRFAM, and HAMAP databases) (87–90), and Clinker was used to generate comparisons of annotated predicted proteins (91).

### Identification of CRISPR arrays in bacterial genomes

MinCED and CRISPRCasDB were used to identify CRISPR repeats in the smaller set of *Neisseria* genomes (92, 93). Using the CRISPR repeats identified in high-quality genomes, we next used a custom script (reps2spacers.py; https://github.com/Alan-Collins/Neisseria-prophage-paper) to run BLASTn (using -task blastn-short) and process results to identify spacers in the larger set of *Neisseria* genomes (81). CRISPRCasTyper was used to predict the CRISPR subtype associated with each of the identified repeats (94). To investigate sharing of identical spacers between genomes of different species, an all-by-all BLASTn (using -task blastn-short) was performed on all spacers that were identified in the larger *Neisseria* genome dataset (81).

### Prediction of CRISPR targeting of prophages and bacterial genomes

BLASTn (using -task blastn-short) was used to identify matches between *Neisseria* CRISPR spacers (identified in the larger *Neisseria* genome dataset) and either prophages or high-quality bacterial genomes (81). Matches were filtered as follows: the spacer had to match the target with 100% identity over the entire length of the spacer. Additionally, matches between spacers and CRISPR arrays found within bacterial genomes or prophages were removed. CRISPRopenDB was used to predict CRISPR targeting of predicted prophages by other bacterial taxa using the default setting of 2 mismatches (65).

### Comparison of CRISPR targeting of prophages vs. bacterial genome backbones and statistical testing

To compare the CRISPR targeting of predicted prophages and the genomes in which they were found (i.e., the backbone), we used the following method. As described above, matches between CRISPR spacers and targets in prophages or bacterial genomes were identified using BLASTn (81), and only hits with 100% identity over the full length of the spacer were kept. CRISPR spacers were excluded if they matched a CRISPR array found in either a predicted prophage or bacterial genome. Next, we quantified CRISPR targeting per kb; importantly, this was done differently for prophages and backbones as follows. For prophages, the targeting density is the number of CRISPR matches divided by the prophage length in kb. For backbones, targeting is the number of matches within the entire bacterial genome minus any matches to locations that are known/predicted to be part of an MGE; the length is calculated by subtracting the length of all MGEs identified in the genome from the length of the entire bacterial genome. Then, the backbone targeting density was calculated as the number of CRISPR targets (not in a known/predicted MGE) divided by the length of the genome minus the lengths of all known/predicted MGEs. Afterwards, we performed statistical testing to test whether there is a difference between the targeting density of the prophage and the backbone for each prophage; to do this, each kb of prophage or bacterial genome was treated as a separate datapoint. Specifically, we performed a Mann-Whitney U test to compare each of the CRISPR targeting counts for the separate kb bins between each prophage and the backbone using the Python package SciPy (82, 83). P-values were adjusted with the Holm-Sidak correction using the Python package Statsmodels (82, 95).

### Inferring host bacterial species of known and predicted prophages

Results from the interspecies CRISPR targeting analysis described above were used to infer the additional host species of known *Neisseria* phages and dereplicated predictions made by PhiSpy, VirSorter2, and Seeker. Any species that was found to target a predicted prophage with ≥1 spacer was inferred to be a host of that predicted prophage. Only dereplicated predicted prophages that were found to have a significantly higher targeting density compared to the rest of the genome in which they were identified (as described above) were included in this analysis.

## Data availability

All of the genome sequences used in this study were downloaded from GenBank. Accession numbers of the 2619 bacterial genome assemblies used in this study are included in **Table S1D**. Accession numbers of phage genomes and plasmids are found in **Table S1B** and **S1C**, respectively. Custom scripts created for analysis of data in this study are available at https://github.com/Alan-Collins/Neisseria-prophage-paper. Key data sets generated in this study and resulting analyses are provided in the Supplemental Material.

## ACKNOWLEDGMENTS

We thank Julie Pryde, Awais Vaid, and other members of the Champaign-Urbana Public Health District for their collaboration, which inspired us to conduct this study.

This work was supported by an Allen Distinguished Investigator award to R.J.W (ADI12345) and the Carl R. Woese Institute for Genomic Biology Postdoctoral Fellowship to G.O.

## SUPPLEMENTAL MATERIAL

**Fig. S1. Comparison of known *Neisseria* prophages to predictions made by three bioinformatic tools**. (A and B) Visualizations of locations of known and predicted prophages made using SnapGene software (from Insightful Science; available at snapgene.com) and modified. PhiSpy, VirSorter2, or Seeker were used to predict prophages in two bacterial genomes: *N. gonorrhoeae* FA10 (A) and *N. meningitidis* Z2491 (B). Known dsDNA prophages are shown in dark grey, filamentous prophages in light grey, and predictions are shown in different colors as indicated. Known *Neisseria* prophages are described in Table S1B, and information about each prediction is provided in Table S2A.

**Fig. S2. The distribution of predicted prophage lengths for each virus prediction tool**. (A to C) Length in kb is shown for dereplicated PhiSpy (A), VirSorter2 (B), and (C) Seeker predictions. Information about each prediction is provided in Table S2A and dereplication of predictions in Table S2B.

**Fig. S3. Few predictions cluster with known *Neisseria* plasmids or genetic island based on percent-length aligned nucleotide sequences**. (A and B) Dendrogram representing average-linkage hierarchical clustering of known *Neisseria* plasmids (A) and the Gonococcal Genetic Island (GGI) (B) with dereplicated predictions from all tools based on percent length aligned by BLASTn. Clustering was performed using SciPy and the resulting dendrograms were visualized with iTOL. Distinct clusters are highlighted by alternating light and dark grey clade shading. Color strips indicate the *Neisseria* species in which the MGE was identified and whether the MGE was previously known or predicted in this study. Beneath the color strips is an identifying cluster number for each cluster. Only predictions that cluster with known *Neisseria* plasmids or the GGI are included in the dendrograms. Information about plasmids and the GGI are provided in Table S1C. Hierarchical clustering memberships of each indicated cluster are presented in Data Set S1, Tab 2.

**Fig. S4. A greater number of PhiSpy and VirSorter2 predictions are connected to reference viruses compared to Seeker predictions**. (A and B) vConTACT v.2.0-generated networks of dereplicated VirSorter2 (A) or Seeker (B) predictions with reference viruses visualized with Cytoscape using an edge-weighted spring-embedded algorithm. Nodes represent reference viruses (dark grey), predicted prophages (color corresponding to the *Neisseria* species in which they were identified), or known *Neisseria* phages (dark grey outlined in the color corresponding to the bacterial host species). Edges show the vConTACT v.2.0-generated similarity score between each pair of viruses (only similarity scores ≥1 are included in the network). Highly similar viruses are positioned close together. Only reference viruses that connect to ≥1 predicted prophage are included in the network.

**Fig. S5. PhiSpy and VirSorter2 predictions are more similar to reference viruses compared to Seeker predictions**. vConTACT v.2.0-generated similarity scores between each predicted prophage and its most similar reference virus in the network. Similarity scores are shown for each virus pair and grouped by the tool that predicted the prophage (PhiSpy, VirSorter2, or Seeker). Distributions of similarity scores were compared between each tool using the Mann-Whitney U test. Similarity scores (edge weights) for each network are provided in Data Set S1, Tabs 4-6.

**Fig. S6. Inferred additional host species of each PhiSpy prediction grouped by vConTACT cluster**. Interspecies CRISPR targeting data were used to infer additional host species for each predicted prophage. The distribution of the number of inferred additional host species is shown for PhiSpy-predicted prophages (only dereplicated predictions that have a significantly higher CRISPR targeting density compared to the bacterial genome backbone; listed in Data Set S1, Tab 11). The data are grouped according to the vConTACT v.2.0 subcluster.

**Table S1**. (A) Smaller set of *Neisseria* genome assemblies used in this study to predict prophages. (B) Prophages previously identified in *Neisseria* genomes. (C) *Neisseria* plasmids and the Gonococcal Genetic Island. (D) Larger set of *Neisseria* genome assemblies used in this study to identify CRISPR arrays.

**Table S2**. (A) Prophages predicted in this study. (B) Dereplication of predicted prophages at 95% length aligned. (C) Comparison of clustering and CRISPR targeting of predicted prophages between tools. (D) Inferred additional *Neisseria* host species of significantly targeted prophages.

**Data Set S1**. Data included in each tab: (1) Bacterial genome maximum-likelihood ribosomal MLST tree file. (2) Predictions that cluster with *Neisseria* plasmids and the Gonococcal Genetic Island by hierarchical clustering. (3) vConTACT subcluster information and memberships. (4-6) Edge weights of vConTACT networks. (7) CRISPR arrays identified in each *Neisseria* genome. (8) *Neisseria* CRISPR repeat sequences and associated subtypes. (9) *Neisseria* CRISPR spacer sequences. (10) CRISPR targeting densities of known and predicted prophages compared to genome backbone targeting density. (11) Dereplicated prophages with CRISPR targeting densities that are significantly higher than the bacterial genome. (12) vConTACT subclusters that include significantly targeted predicted prophages. (13) Identical spacers identified in genomes of different *Neisseria* species. (14) Matches between spacers encoded by other bacterial taxa and *Neisseria* predicted prophages.

## Notes

### Competing Interest Statement

The authors have declared no competing interest.

### Summary of Updates

Figures 7 and 8 were revised.

